# Accelerating structure-function mapping using the ViVa webtool to mine natural variation

**DOI:** 10.1101/488395

**Authors:** Morgan O. Hamm, Britney L. Moss, Alexander R. Leydon, Hardik P. Gala, Amy Lanctot, Román Ramos, Hannah Klaeser, Andrew C. Lemmex, Mollye L. Zahler, Jennifer L. Nemhauser, R. Clay Wright

## Abstract

Thousands of sequenced genomes are now publicly available capturing a significant amount of natural variation within plant species; yet, much of this data remains inaccessible to researchers without significant bioinformatics experience. Here, we present a webtool called ViVa (Visualizing Variation) which aims to empower any researcher to take advantage of the amazing genetic resource collected in the *Arabidopsis thaliana* 1001 Genomes Project (http://1001genomes.org). ViVa facilitates data mining on the gene, gene family or gene network level. To test the utility and accessibility of ViVa, we assembled a team with a range of expertise within biology and bioinformatics to analyze the natural variation within the well-studied nuclear auxin signaling pathway. Our analysis has provided further confirmation of existing knowledge and has also helped generate new hypotheses regarding this well studied pathway. These results highlight how natural variation could be used to generate and test hypotheses about less studied gene families and networks, especially when paired with biochemical and genetic characterization. ViVa is also readily extensible to databases of interspecific genetic variation in plants as well as other organisms, such as the 3,000 Rice Genomes Project (http://snp-seek.irri.org/) and human genetic variation (https://www.ncbi.nlm.nih.gov/clinvar/).

## Introduction

The sequencing of the first *Arabidopsis thaliana* genome ushered in a new era of tool development and systematic functional annotation of plant genes (The Arabidopsis Genome Initiative 2000). Since that landmark effort, massive scaling of sequencing technology has allowed for the survey of genomic variation in natural *Arabidopsis thaliana* populations (Nordborg et al. 2005; Borevitz et al. 2007; Weigel and Mott 2009). This valuable population genetics resource has led to several associations of genetic loci with phenotypic traits and provided insights into how selective pressure has influenced the evolution of plant genomes (Long et al. 2013; Atwell et al. 2010; Clark et al. 2007).

Beyond its utility in gene discovery and understanding genome evolution, natural genetic variation provides a catalog of permissible polymorphisms that can facilitate the connection of genotype to phenotype at the gene, gene family and network scales (Joly-Lopez, Flowers, and Purugganan 2016). This is an especially critical resource in large gene families where loss of function in individual genes may have little or no phenotypic effect and directed allele replacement remains time and resource-intensive. Massively parallel assays of variant effects in human clinical medicine stand to revolutionize genetic diagnostics and personalized medicine (Starita et al. 2017; Gasperini, Starita, and Shendure 2016; Matreyek, Stephany, and Fowler 2017). We envision the use of plant natural variation datasets as a tool to similarly revolutionize breeding and genetic engineering of crop plants by rapidly advancing our understanding of genotype/function/phenotype relationships. A proof-of-principle survey of a relatively small subset of natural variants paired with a synthetic assay of gene function successfully mapped critical functional domains of auxin receptors and identified new alleles which affect plant phenotype (Wright et al. 2017).

Why is the survey of natural variants not as routine as a BLAST search or ordering T-DNA insertion mutants? One reason may be the current requirement for a fairly high level of bioinformatics expertise to extract the desired information from whole genome resequencing datasets. While existing resources such as the 1001 Proteomes website (Joshi et al. 2012) and ePlant (Waese et al. 2017) facilitate access to this data at the gene scale, they do not provide summaries or visualizations of variation at the gene family and network scales. To address this concern, we created ViVa: a webtool and R-package for **Vi**sualizing **Va**riation, which allows plant molecular biologists of any level access to gene-level data from the 1001 Genomes database. Using ViVa researchers may: 1) Identify polymorphisms to facilitate biochemical assays of variant effects (Starita et al. 2017; Wright et al. 2017); 2) Produce family-wise alignments of variants to facilitate *de novo* functional domain identification (Melamed et al. 2015); 3) Generate lists of accessions containing polymorphisms to facilitate phenotypic analysis of gene variant effects (Park et al. 2017); and 4) Quantify metrics of genetic diversity to facilitate the study of gene, gene family and network evolution (Delker et al. 2010; Kliebenstein 2008).

## Results and Discussion

### An overview of ViVa

ViVa, in this first iteration, is meant to visualize natural variation in the coding sequences of genes. Non-coding sequence variation is intentionally excluded from the analysis tools. This reflects challenges both in alignment of non-coding sequences and the increased difficulty in assessing variation in these regions (Alexandre et al. 2018).

The development version of ViVa can be accessed as a Docker container https://hub.docker.com/r/wrightrc/r1001genomes/ or as an R-package at https://github.com/wrightrc/r1001genomes. ViVa will be hosted at https://www.plantsynbiolab.bse.vt.edu/ViVa/ upon release.

**Figure 1.**
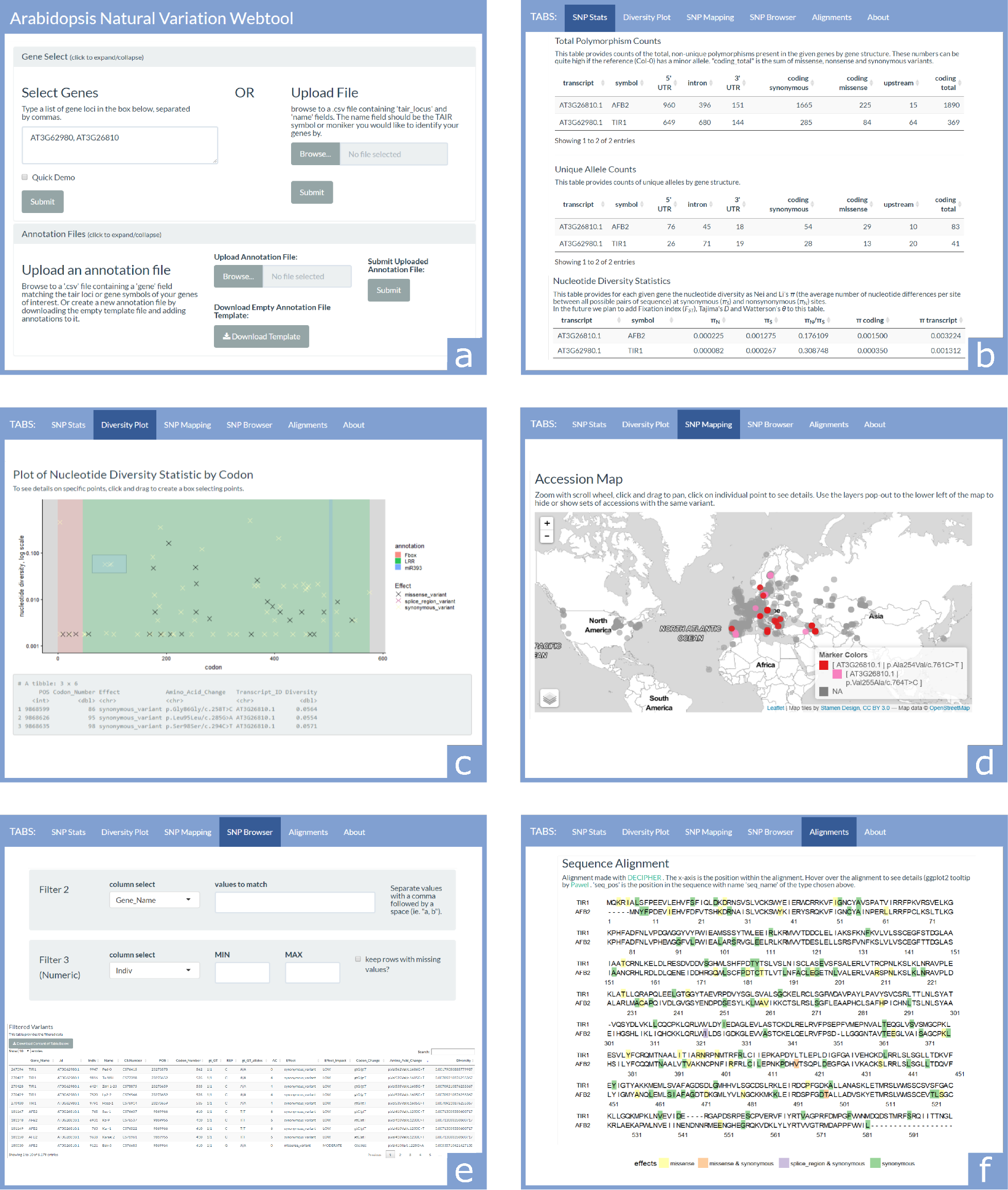
Key elements of the webtool. (a) The first section contains two collapsible panels, Gene Select and Annotation Files, which are used to input information about the genes to be investigated. (b) The SNP Stats tab provides gene-structure level counts and statistics on SNPs. (c) The Diversity Plot tab plots the nucleotide diversity of SNP sites along the length of the coding region of a selected gene. (d) The SNP Mapping tab plots accessions on a world map colored according to the selected set of SNPs. (e) The SNP Browser tab allows variants and accessions to be filtered by any combination of text and numeric fields. (f) The Alignments tab aligns DNA and amino acid sequences of homologous genes, and colors sequence elements based on SNPs and annotations.

### Gene Select and Annotation Files

At the top of the ViVa webtool are two collapsible panels used for entering the genes to query and custom annotations for those genes (Figure 1a). The Gene Select panel permits gene input by either typing in the genes’ AGI/TAIR locus identifiers, or uploading a.csv file. The Annotation Files panel is optionally used to upload an annotation file containing coordinates of domains, mutations, or any other sequence knowledge that will be plotted on some of the tabs of the webtools analysis section.

Below the data input section, the rest of the webtool is divided into several tabs containing the interactive output of the program.

### SNP Stats: Summary of gene information, structure, and diversity

The SNP Stats tab provides general information on the gene transcripts being queried, as well as calculated counts/statistics on the content of variants found in the sample population (Figure 1b). The first table of this section is basic information about the transcripts, including TAIR locus and symbol, the chromosomal start and end position and the transcript length. This information was collected from the Araport11 Official Release (06/2016) annotation dataset (Cheng et al. 2017).

The next two tables provide counts of SNPs across the gene body for each transcript. The Total Polymorphism Counts table provides the total number of observations of non-reference allele counts of each variant type (the Col-0 accession TAIR9 genome is the reference genome for this dataset). The Unique Allele Count table only counts the number of unique variants and alleles within the population of accessions (*e.g*. if multiple accessions have the same variant allele, that allele will only be counted once).

The Nucleotide Diversity Statistics table provides a nucleotide diversity statistic (*π*) for the transcript and the coding sequence of each gene (Nei and Li 1979). Nucleotide diversity is also calculated for the set of only synonymous (*π*_*S*_) and only non-synonymous sites (*π*_*N*_). The ratio of the presence of non-synonymous to synonymous polymorphism provides a measure of the potential for functional diversity (Firnberg and Ostermeier 2013; Whitehead et al. 2012). We present *π*_*N*_/*π*_*S*_ here as an correlate for functional diversity throughout ViVa (Nelson, Moncla, and Hughes 2015; Hughes et al. 2000). While imperfect, this metric may be suggestive of functional constraint when *π*_*N*_/*π*_*S*_ ≪ 1 and functional diversity when *π*_*N*_/*π*_*S*_ ≫ 1 (Hughes 1999).

### Diversity plot: Visualize allelic diversity across the coding sequence

The Diversity Plot tab shows the nucleotide diversity of each variant in the coding region of a selected gene (Figure 1c). Although the X-axis is marked by codon number from the N-terminus for interpretability, the diversity values are based on single nucleotide sites. The colors of markers on the plot identify the effect of the polymorphism. If annotation files are provided, the background of the plot will be color coded by the annotated regions. If points on the plot are selected by clicking and dragging a box over them, the data for the selected points will appear in the grey box below the plot. Below this is a complete data table containing all points on the plot which can be downloaded as a.csv file. This tab allows users to identify regions of high diversity as well as isolate polymorphisms that may affect gene function and exist in multiple accessions, facilitating phenotypic analysis.

### SNP Mapping: View distributions of SNPs across the globe

The SNP Mapping tab plots the accessions collection locations on a world map, and colors the points based on selected variant alleles (Figure 1d). After selecting the genes and filtering on the SNP type and level of nucleotide diversity, a group of checkboxes becomes available to select variant alleles to display on the map. The variant alleles are labeled with the Transcript_ID and Amino_Acid_Change fields, in the form [Transcript_ID|Amino_Acid_Change]. After selecting the variant alleles and updating the map, the accessions on the map will be colored by each present unique combination of the selected alleles. Below the map is a table containing the accession details for all mapped accessions. This tab may help users formulate hypotheses about the relatedness of accessions sharing a common allele and environments in which that allele may be favorable.

### SNP Browser: Filter and search for variants

The SNP Browser tab provides a way to search and filter the variant data by different fields (Figure 1e). After selecting the transcripts to include, a number of filters can be applied to the dataset to match text values (e.g. gene name or variant effect), or set minimum and maximum limits on the values of numeric fields (e.g. nucleotide diversity). When these filters are applied, the table below will update to only contain rows meeting the criteria for all filters. This tab can be useful for identifying all accessions with a particular SNP allele, or any non-reference alleles in a particular region of a gene that may not have been easily accessible in another tab.

### Alignments: Visualize SNPs on alignments of homologous genes

The Alignments tab provides DNA and amino acid sequence alignments of selected genes, colored according to the variant allele with the strongest functional effect at each position (Figure 1f and 2). The content of this tab is most useful if the selected genes are all family members or have significant sequence homology. If annotation files are uploaded, the background of the sequences will be colored by annotation region. Hovering the cursor over variants will provide additional details about the alleles present at that locus. This tab facilitates family-wise analysis of functional conservation, allowing users to identify potential functional regions and alleles which may be useful in deciphering this function.

### Gene Tree: Visualize functional diversity and sequence divergence of a gene family

The Gene Tree tab provides a neighbor joining tree (or uploaded tree created by the user) for the selected genes with the tips of the tree mapped with predicted functional diversity as represented by *π*_*N*_/*π*_*S*_ in the 1001 Genomes dataset (Figure 3). This tab allows users to generate hypotheses regarding functional diversity and redundancy within the context of the predicted evolution of the gene family.

### ViVa R package: Programmatic access to ViVa’s functionality

All of the functionalities of the ViVa webtool are implemented through function calls to the ViVa R package. In addition to being able to generate the same sets of figures and tables as in the webtool, users of the R package also gain direct access to the underlying data structures, providing greater control over parameters when processing and visualizing the data. The ViVa R package can be found at https://github.com/wrightrc/r1001genomes and can be installed in your R environment via the devtools package: `devtools::install_github(“wrightrc/r1001genomes“)`.

### Visualizing Variation within the auxin signaling pathway

To test the usability and accessibility of ViVa, we assembled a group of alpha testers comprising postdoctoral, graduate, and undergraduate researchers at a research university (University of Washington) and at a primarily-undergraduate institution (Whitman College). Our testers focused their investigation of natural variation on the nuclear auxin signaling pathway. We selected this signaling pathway for multiple reasons including a wealth of functional data and solved structures of several domains or entire proteins. Using this exitsting knowledge, we were able to qualitatively assess the predictive ability of the ViVa modules. Summary information about each nuclear auxin signaling gene family examined can be found in the Supplemental Data. Below, we describe the results for the *Aux*/*IAA* family in more detail.

In most cases, natural selection is expected to minimize the persistence of nonsynonymous mutations in sequences that encode critical functional domains relative to their persistence in non-critical domains (Hughes et al. 2000). Therefore, we reasoned that scanning gene coding sequences for regions of relatively low nonsynonymous diversity should highlight functional domains. This general principle can be seen clearly in the analysis of the *Aux*/*IAA* family of transcriptional co-repressors/co-receptors. Aux/IAAs have three major domains. Domain I contains an EAR motif that facilitates interaction with TOPLESS (TPL) and TOPLESS-related (TPR) transcriptional repressors (Tiwari, Hagen, and Guilfoyle 2004; Szemenyei, Hannon, and Long 2008). Domain II, the degron, facilitates interactions with the TIR1/AFB receptors in the presence of auxin (Tan et al. 2007). Domain III (which was originally considered domains III and IV) is a PB1 domain and facilitates interactions with the ARF transcription factors (Ulmasov et al. 1997; Guilfoyle and Hagen 2012; Nanao et al. 2014; Korasick et al. 2014). The EAR motif and degron can be readily identified by the drop in nonsynonymous variation, as visualized by the lack of stronger functional effects in Figure 2. The PB1 domain is not as readily identified, perhaps because the multiple contributing residues are spread out in linear sequence space. It is worth noting that the charged residues that facilitate electrostatic PB1-PB1 interaction show little variation.

**Figure 2.**
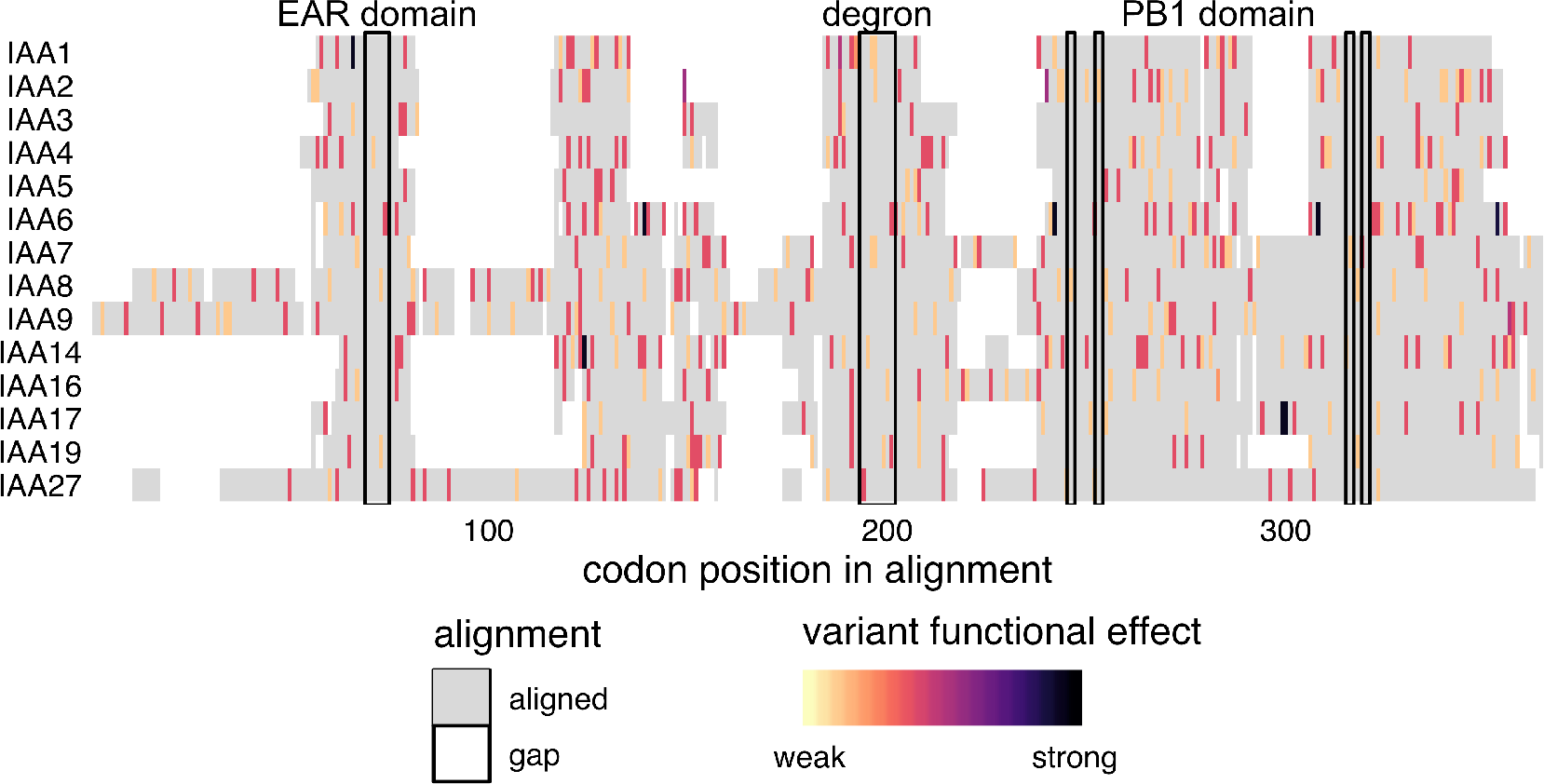
Canonical Aux/IAAs have conserved Degron and EAR motifs. Protein sequences were aligned with DECIPHER (Wright 2015) and variants were mapped to this alignment and colored according to the predicted functional effect of the allele of strongest effect at that position, with light colors having weaker effects on function and darker colors stronger effects. Red indicates missense variants. Color scale is explained in Methods.

Natural variation also provides a means to study how gene families are evolving. To do this, we used ViVa to map onto the Aux/IAA phylogenetic tree the diversity at nonsynonymous variant sites relative to synonymous sites (Figure 3). This visualization enables straightforward comparison of rates of recent functional divergence within the context of rates of sequence divergence across the entire gene family. By comparing nonsynonymous diversity gene clades can be identified that are likely to exhibit high rates of functional conservation and possible redundancy or conversely, where there is a possibility for recent emergent novel function or pseudogenization.

Previous research has found evidence of both broad genetic redundancy among the *Aux*/*IAAs* and also specificity within closely related pairs or groups of Aux/IAA proteins (Overvoorde et al. 2005; Winkler et al. 2017). For example, the *iaa8-1 iaa9-1* double mutant and the *iaa5-1 iaa6-1 iaa19-1* triple mutant have wild-type phenotypes (Overvoorde et al. 2005), yet the *IAA6*/*IAA19* sister pair has significant differences in expression patterns, protein abundances and functions suggesting they have undergone functional specialization since their divergence (Winkler et al. 2017). A closer examination of the *IAA19* and *IAA6* pair within *Brassicaceae* found evidence for positive selection and possible subfunctionalization of *IAA6* relative to *IAA19* (Winkler et al. 2017). Consistent with these results, ViVa revealed higher conservation for *IAA19* (*π*_*N*_/*π*_*S*_ = 0.55) compared to *IAA6* (*π*_*N*_/*π*_*S*_ = 2.3) (Figure 3), and also detected high nonsynonymous diversity within the same regions of *IAA6* as seen by Winkler et al. (Figure 8). This pattern—one sister showing high nonsynonymous diversity while the other sister was more conserved—was observed frequently across the *Aux*/*IAA* as well as the *AFB* and *ARF* families (Figure 5 and 11), suggesting this could be a recurring feature in the evolution of these families supporting the large diversity in auxin functions.

**Figure 3.**
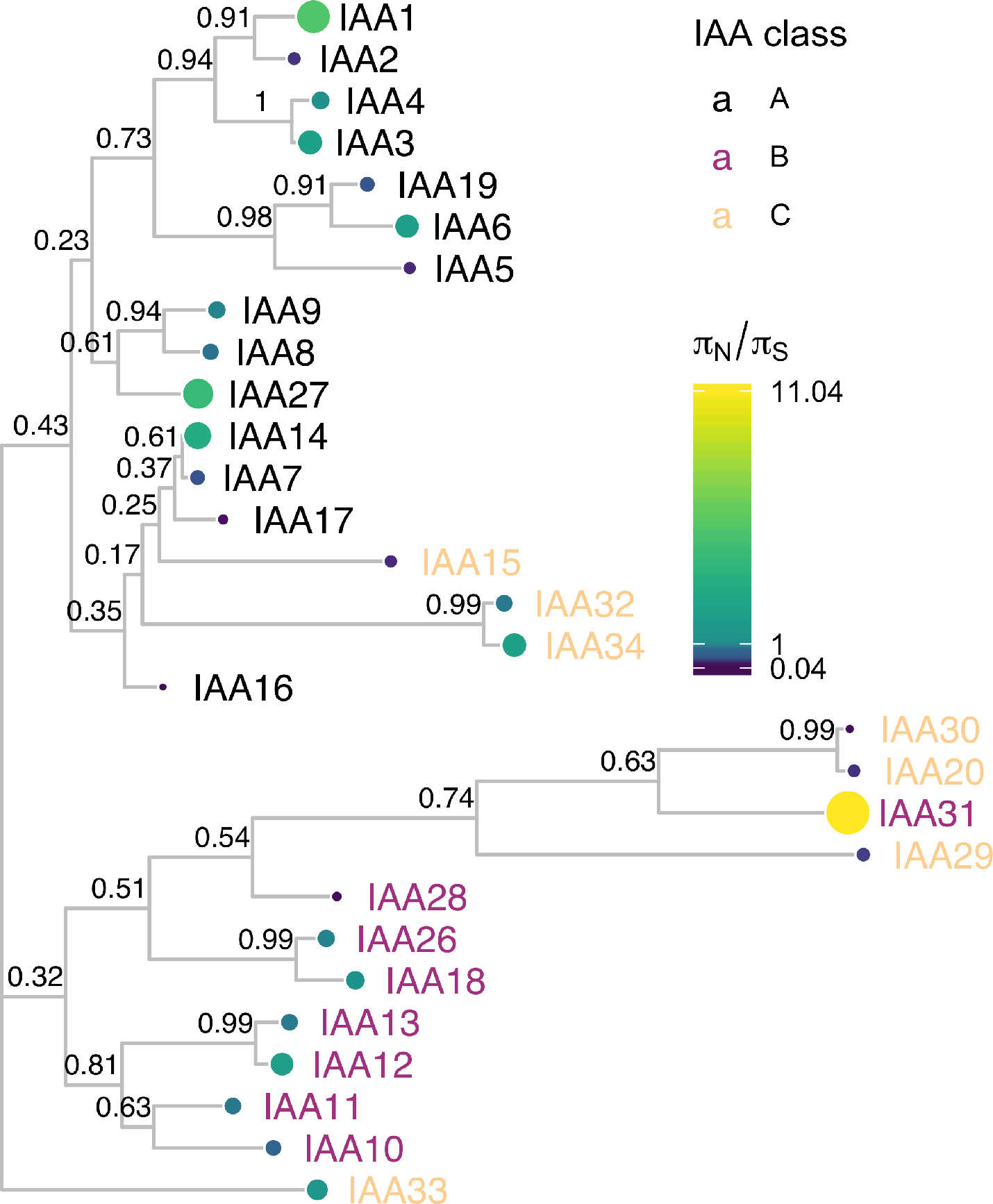
IAA protein sequence tree mapped with *π*_*N*_/*π*_*S*_. Protein sequences were aligned with DECIPHER (Wright 2015) and low information content regions were masked (Kück et al. 2010) prior to inferring a phylogeny (Ronquist and Huelsenbeck 2003). Tips of the tree are mapped with circles of color and diameter proportional to *π*_*N*_/*π*_*S*_. *π*_*N*_/*π*_*S*_ provides a prediction of functional diversity. Nodes are labeled with the poster probability of monophyly. There are two distinct clades of Aux/IAAs represented by the majority of the A and B classes. C class Aux/IAAs are missing one or more of the canonical Aux/IAA domains

The *Aux*/*IAA* phylogeny clusters into two distinct clades represented by the A and B classes (Remington et al. 2004). The C class Aux/IAAs are missing one or more of the canonical Aux/IAA domains. We found notable exceptions to pattern of diversification and conservation between sister pairs within the Class B *Aux*/*IAA* genes. The *IAA10*/*IAA11*, *IAA18*/*IAA26*, and *IAA20*/*IAA30* pairs showed similar levels of nonsynonymous diversity. For example, *IAA10* and *IAA11* both showed functional conservation (*π*_*N*_/*π*_*S*_ of 0.80 and 0.67 respectively). In support of this strong conservation of both *IAA10* and *IAA11*, the *Arabidopsis thaliana* ePlant browser indicates that *IAA10* and *IAA11* have almost identical expression patterns (Waese et al. 2017). Together this evidence suggests a strong dosage requirement for these genes or that they have taken on novel functions since their emergence.

## Conclusion

ViVa has allowed our team of testers from various skill levels and backgrounds to meaningfully access the 1001 genomes dataset. The visualizations of natural variation further supported much of the existing structure-function knowledge of this well studied signaling pathway and facilitated the generation of new hypotheses. Application of ViVa to less studied genes and gene families promises to yield more novel hypotheses, which can be evaluated with mutagenesis and functional assays to glean novel structure/function knowledge from this rich dataset.

ViVa results are intended to inform and inspire hypothesis generation, not be taken as absolute evidence of trends in gene or gene family evolution. Among the cautions worth noting in interpreting results are limitations of short-read sequencing that lead to regions of missing data where low read quality may have prevented variant calls. We have assumed these missing variants are reference alleles, leading to undercounting in ViVa’s diversity estimations. Recent advances in sequencing technologies have been combined to generate extremely high quality genomes (Michael et al. 2018), and will reduce this source of uncertainty in future resequencing datasets. Another limitation is that the geographic coverage of accessions in the 1001 Genomes dataset is far from uniform, and thus diversity scores may not accurately reflect the allelic distributions of the global *Arabidopsis thaliana* population.

We hope that ViVa will advance understanding of genotype-phenotype relationships by allowing all researchers access to large resequencing datasets. In the future, we intend to expand ViVa beyond the plant genetics workhorse, *Arabidopsis thaliana*, to more agriculturally relevant species with existing resequencing projects, such as rice (Wang et al. 2018) and soybean (Zhou et al. 2015). Indeed, the ViVa framework is readily adaptable to any source of targeted resequencing data. If François Jacob’s metaphor holds true, and evolution is indeed a tinkerer and not an engineer (Jacob 1977), it is only by examining the largest possible number of nature’s solutions that we may eventually decipher the principles constraining innovations in form and function.

## Methods

### Data Sources

#### Variant Data

Variant data was queried from the 1001 genomes project (http://1001genomes.org) via URL requests to their API service (http://tools.1001genomes.org/api/index.html). These queries returned subsets of the whole-genome variant call format (VCF) file as SnpEFF VCF files. The whole-genome VCF file can be found on the project’s website at http://1001genomes.org/data/GMI-MPI/releases/v3.1/.

#### Germplasm Accession Information

A dataset of each of the 1135 accessions including CS stock numbers and geographic location where the samples were collected was retrieved from the 1001 Genomes website at http://1001genomes.org/accessions.html, via the download link at the bottom of the page. This data file has been embedded in the R package as **`accessions`**.

#### Gene and Transcript Accession Information

Information on the genes and transcripts including chromosomal coordinates, start and end location, and transcript length were pulled from Araport11 (Cheng et al. 2017). The TAIR10 database, found at http://arabidopsis.org, was accessed via the biomart function, using the biomaRt R-package. The Araport11 full genome general feature format file, which can also be found on the TAIR website (https://www.arabidopsis.org/download/index-auto.jsp?dir=%2Fdownload_files%2FGenes%2FAraport11_genome_release), has been embedded in the R-package as GRanges object, gr. Gene identifiers used in this study are in Table 1.

**Table 1.**
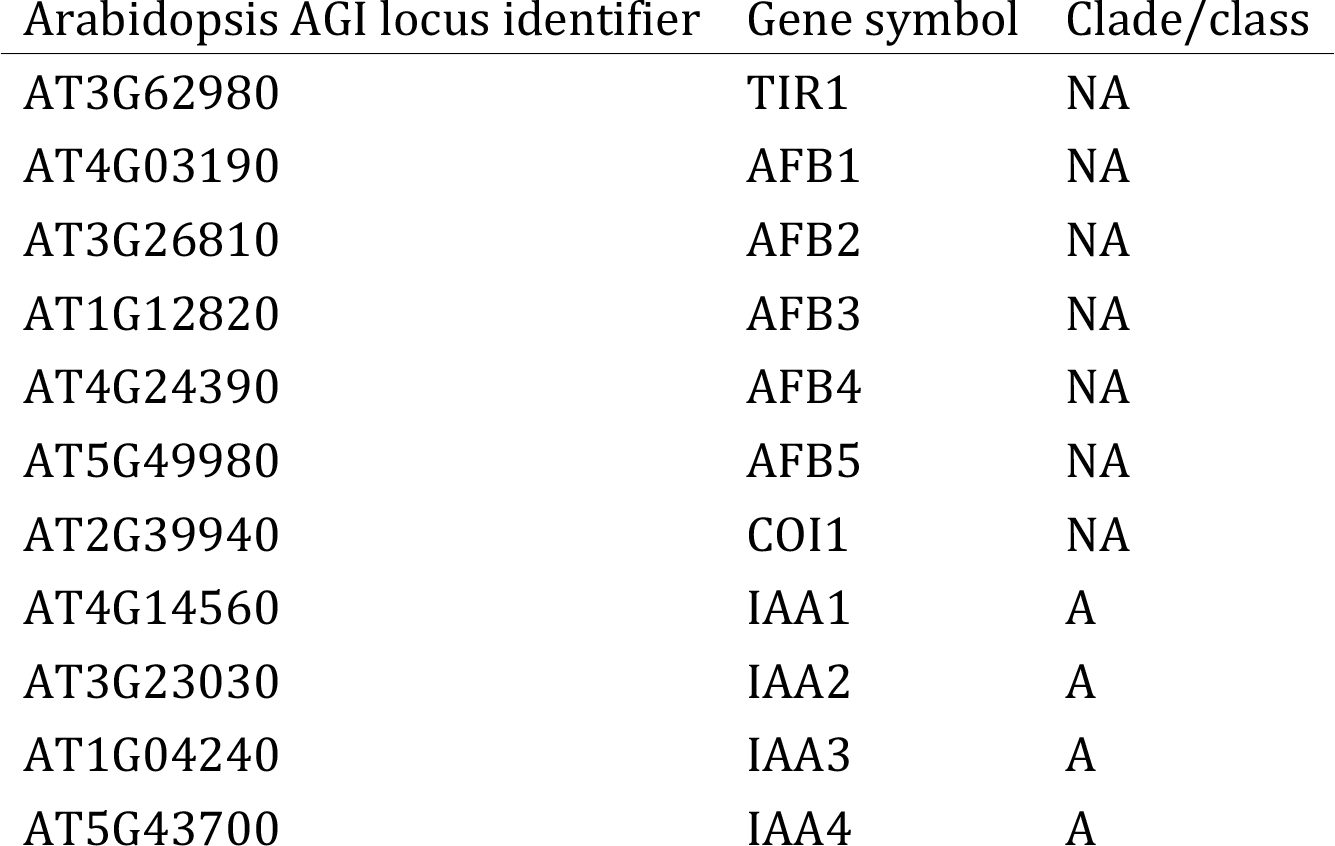
*Full list of genes used in this study, by identifier, symbol and classification*.

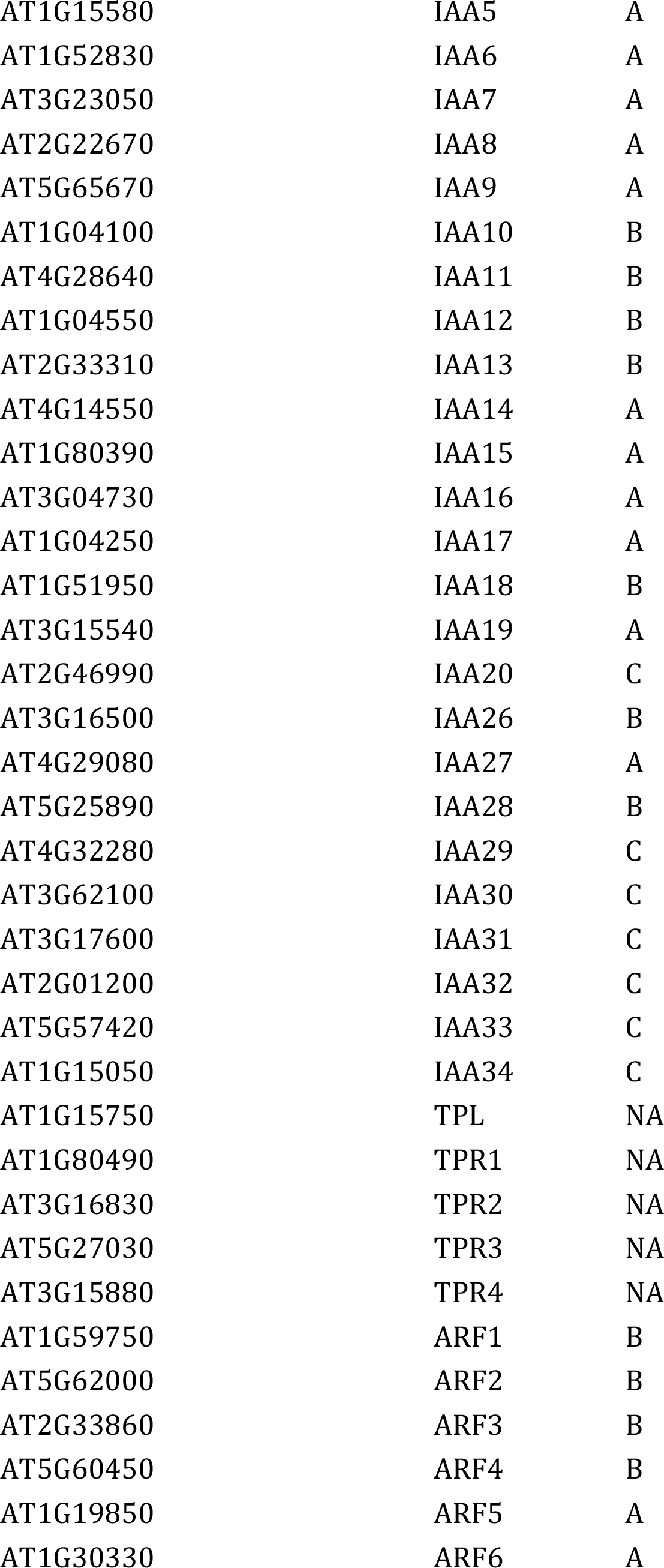

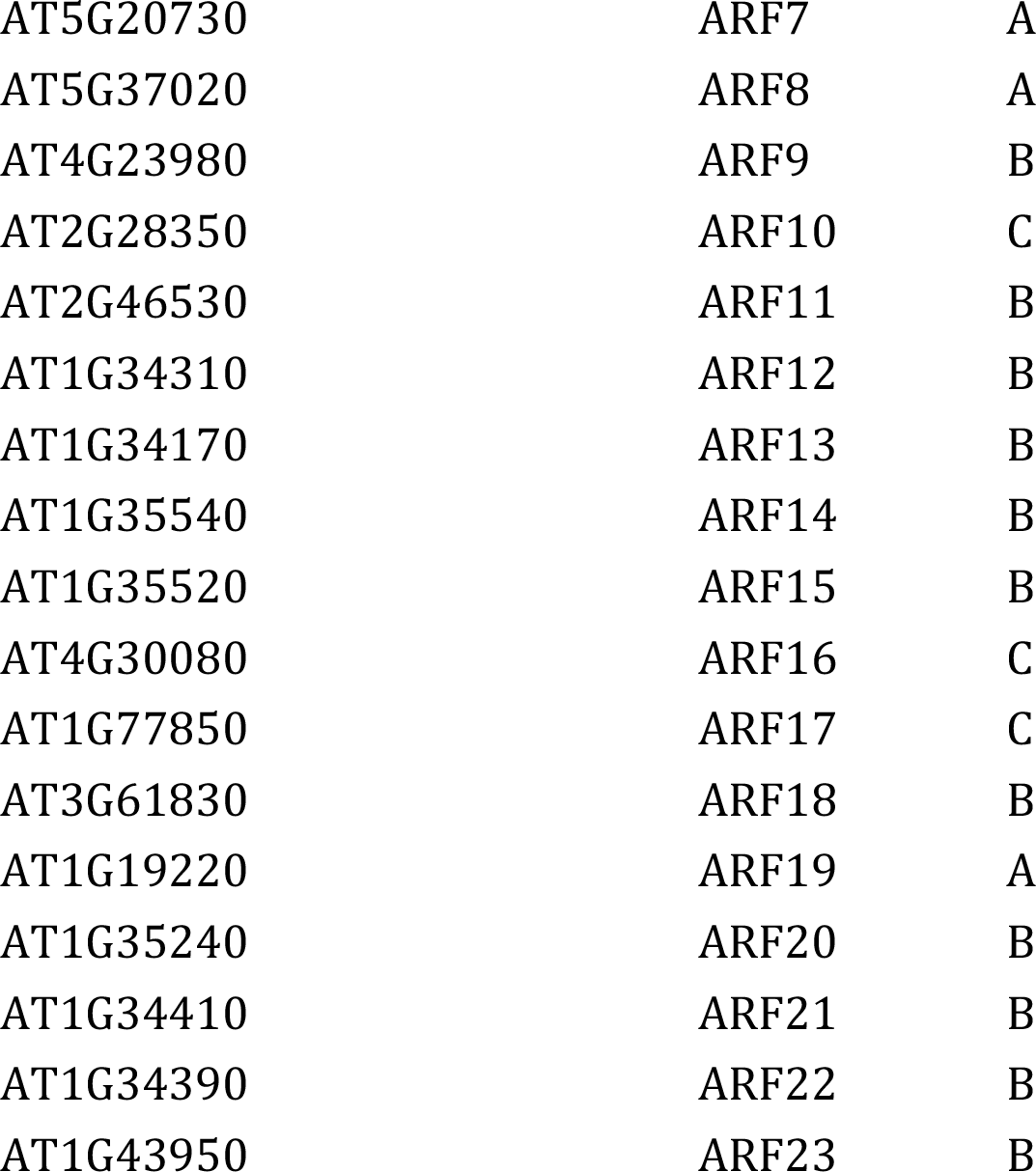

#### Ranking of variant functional effects

Alignments were colored according to the strongest effect variant allele occurring at any frequency at that position as reported in the SnpEFF “effect” field, per the scale in Figure 4.

**Figure 4.**
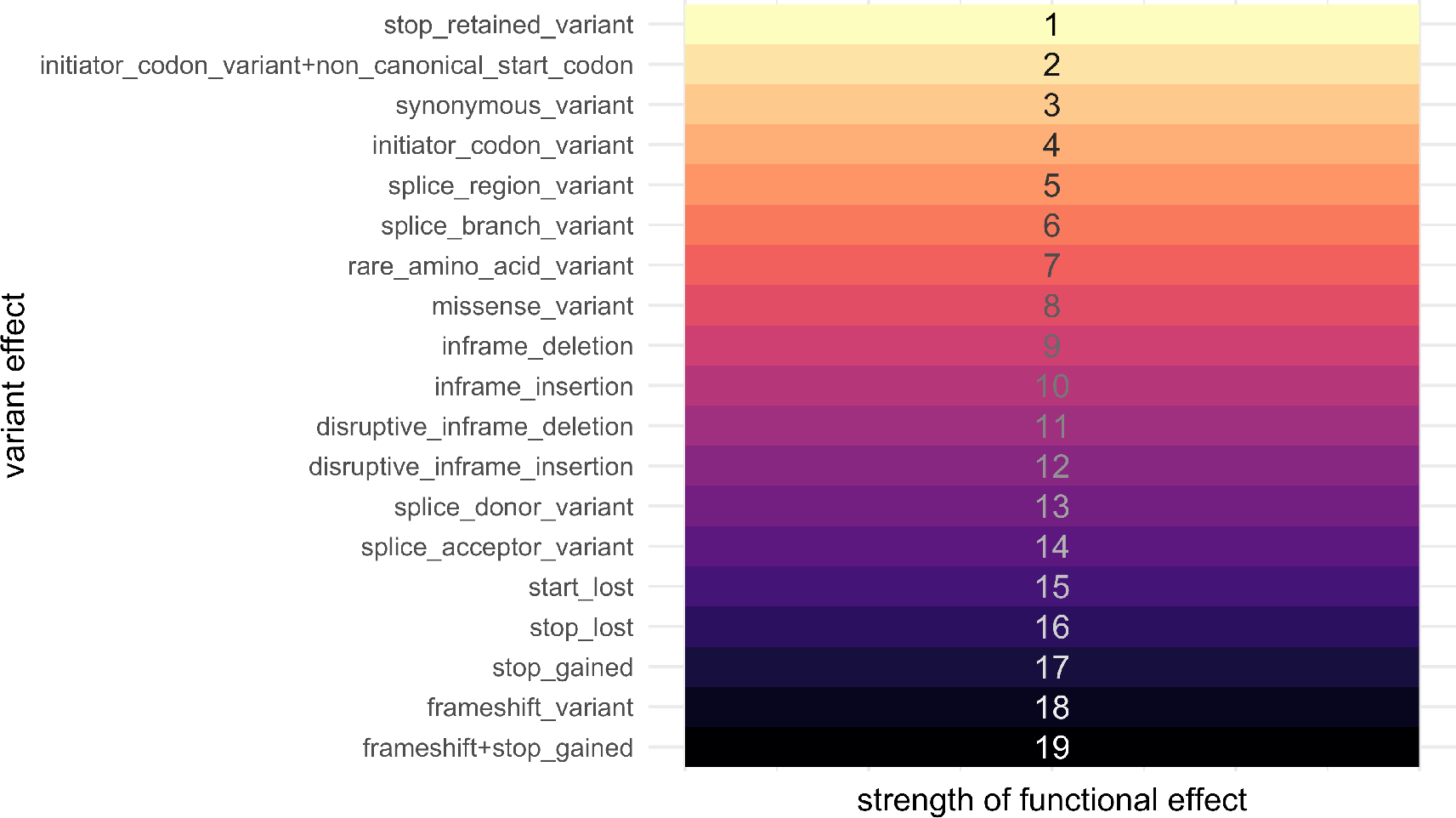
Rank order of the strength of functional effects. Effect classes were ordered by subjective prediction of average strength of effect on gene function. Strength was then assigned to each effect on an integer scale.

### Nucleotide Diversity Calculation

Nei and Li defined the nucleotide diversity statistic in their original paper as: “the average number of nucleotide differences per site between two randomly chosen DNA sequences” (Nei and Li 1979), and provided the equation:

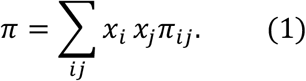

Where *x*_*i*_ is the frequency of the ith sequence in the population and *π*_*ij*_ is the number of sites that are different between the *i* th and; *j*th sequence divided by sequence length.

A more general form, that treats each sequence in the population as unique can be written as:

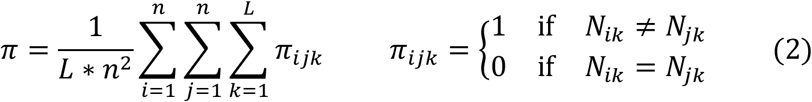

where *N*_*ik*_ is the nucleotide (A, T, C or G) at position *k* on the *i*th sequence of the population. *L* is the length of the sequence. Indels are excluded from the diversity calculation leading to a single *L* for the population. *n* is the total number of sequences in the population.

From this form we can re-arrange summations to the form below:

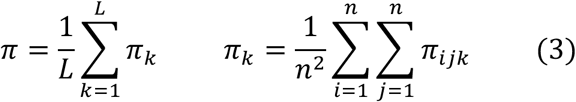

where *π*_*k*_ can be thought of as the site-wise nucleotide diversity at position *k*, and is equal to the nucleotide diversity of a sequence of length 1 at location *k*. We can calculate *π*_*k*_ for each site, then average those over the sequence length to calculate *π*, the nucleotide diversity of the sequence.

The function `Nucleotide_diversity` in the r1001genomes package calculates *π*_*k*_ for each position in the gene or region that contains a variant. Note, *π*_*k*_ is equal to 0 at all locations without variants. This is also what is displayed in the Diversity Plot tab of the webtool.

### Detailed *π*_*k*_ calculation simplification

The formula for *π*_*k*_ above requires comparing every sequence to every other sequence at location k, however, we know there are only a few variant forms at each individual location.

So, we can revert back to using Nei and Li’s original formula (1), modifying it slightly, replacing *x*_*i*_ with 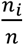, *n*_*i*_ being the number of sequences in the population with nucleotide *N*_*i*_ at location *k*:

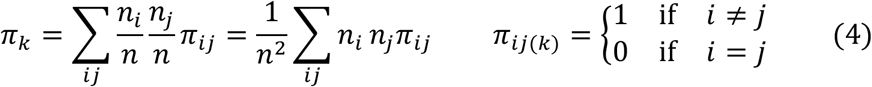

Note that in equation (1) subscripts *i* and *j* are summed over all sequences in the population, however in equation (4) *i* and *j* are only summed over unique variants at a particular location k.

We will define *n*_!*i*_ = *n* -*n*_*i*_ as the number of sequences different from *i* at position *k*. We can also see that the summed term will be zero if *i* = *j*, and *n*_*i*_*n*_*j*_. if *i* ≠ *j*. Therefore:

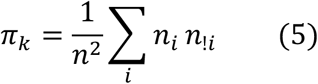

Next we substitute our definition of *n*_!*i*_:

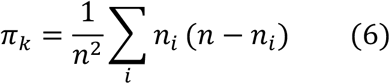

Distributing and splitting summation yields:

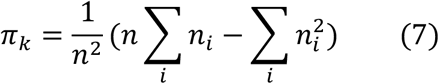

Finally, summing *∑*_*i*_*n*_*i*_ is equal to *n*:

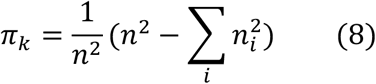

This simplified form for *π*_*k*_ is used by the app, because the counts of unique variants at a single nucleotide location can easily be summarized in R.

### Software

The r1001genomes package has many software dependencies on other R packages, a few of the key bioinformatics packages used are listed below.

#### biomaRt

used for accessing the TAIR10 database on arabidopsis.org

#### vcfR

used to read in the VCF files in a flat “tidy” format for easy manipulation

#### BSgenome

used as the source for the complete DNA string of the reference genome (Col-0).

#### DECIPHER

used to align nucleotide and amino acid sequences of homologous genes

#### GenomicFeatures

used for handling sequence annotations.

#### Biostrings

provides the underlying framework for the sequence manipulations used for generating and aligning sequences with BSgenome, Decipher, and GenomicFeatures

Other packages that were critical to building ViVa and/or writing this document include: (Paradis et al. 2018; R Core Team 2018; Müller 2018; Team 2018; Pagès et al. 2018; Xie 2018a; Ihaka et al. 2016; Wright 2018; Wickham, François, et al. 2018; Xie 2018b; Wickham 2018a; Aphalo 2018a; Wickham, Chang, et al. 2018; Aphalo 2018b; Wagih 2017; Arnold 2018; Yu and Lam 2018; Heibl 2014; Pagès, Aboyoun, and Lawrence 2018; Xie 2018c; Bache and Wickham 2014; Wickham 2016; Henry and Wickham 2018; Hamm and Wright 2018; Neuwirth 2014; Wickham, Hester, and Francois 2017; Wickham 2017a; Allaire, Ushey, and Tang 2018; Allaire et al. 2018; Müller et al. 2018; Pagès, Lawrence, and Aboyoun 2018; Wickham 2018b; Wickham 2018c; Müller and Wickham 2018; Wickham and Henry 2018; Wickham 2017b; Yu 2018; Garnier 2018a; Garnier 2018b; Temple Lang and CRAN Team 2018; Pagès and Aboyoun 2018)

## Acknowledgements

The authors would like to thank Oghenemega Okolo for assistance testing the ViVa software, and Song Li and Bo Zhang for helpful comments on the manuscript. This work was supported by the National Institute of Health (R01-GM107084), the National Science Foundation (IOS-1546873) and the Howard Hughes Medical Institute. R.C.W. received fellowship support from the National Science Foundation (DBI-1402222). B.L.M. and H.K. received support from the M.J. Murdock Charitable Trust. A.R.L. is a Simons Foundation Fellow of the Life Sciences Research Foundation. A.L. was supported by an NSF Graduate Research Fellowship DGE-1256082.

## Supplemental Data

### TIR1/AFB genes

Auxin acts by binding to receptors (Auxin-signaling F-Boxes, or AFBs) that in turn target co-repressors (Aux/IAAs) for degradation. The six auxin receptor genes in the model plant *Arabidopsis thaliana, TIR1* and *AFB1-5*, evolved through gene duplication and diversification early in the history of vascular plants (Parry et al. 2009). The rate of corepressor degradation is determined by the identity of both the receptor and co-repressor (Havens et al. 2012), and this rate sets the pace of lateral root development (Guseman et al. 2015).

All members of this family have been shown to bind auxin and Aux/IAA proteins. However, AFB1 has drastically reduced ability to assemble into an SCF complex, due to the substitution E8K in its F-box domain, preventing it from inducing degradation of Aux/IAAs (Yu et al. 2015). This lack of SCF formation may allow for the high and ubiquitous AFB1 accumulation observed in *Arabidopsis* tissues (Parry et al. 2009). Higher order receptor mutants in the family containing *afb1* mutants suggest that *AFB1* has a moderate positive effect on auxin signaling (Dharmasiri et al. 2005). Additionally, AFB4 and AFB5 have been shown to preferentially and functionally bind the synthetic auxin picloram, while other family members preferentially bind indole-3-acetic acid (Prigge et al. 2016). Interestingly, the strength and rate with which TIR1/AFBs are able to bind and mark Aux/IAAs for degradation are variable (Calderón Villalobos et al. 2012; Havens et al. 2012). AFB2 induces the degradation of certain Aux/IAA proteins at a faster rate than TIR1, suggesting some functional specificity has arisen since the initial duplication between the *TIR1*/*AFB1* and *AFB2*/*AFB3* clades.

**Figure 5.**
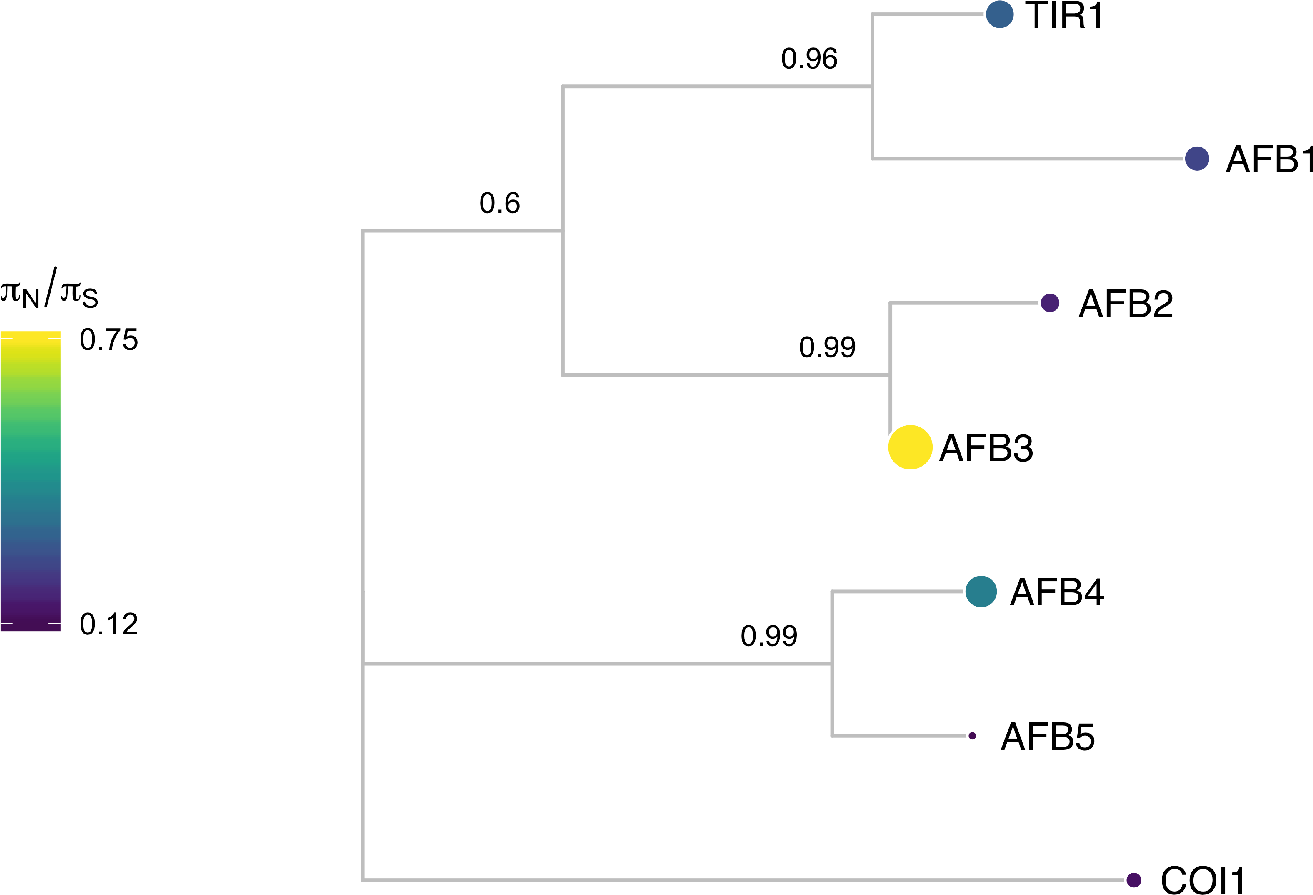
AFB protein sequence tree mapped with *π*_*N*_/*π*_*S*_. Protein sequences were aligned with DECIPHER (Wright 2015) and low information content regions were masked with Aliscore (Kück et al. 2010) prior to inferring a phylogeny with MrBayes (Ronquist and Huelsenbeck 2003). Tips of the tree are mapped with circles of diameter proportional to *π*_*N*_/*π*_*S*_ and also are colored according to *π*_*N*_/*π*_*S*_. Nodes are labeled with the posterior probability of monophyly.

Examining the natural sequence variation across the *AFB* family 5 revealed that *TIR1* and *AFB1* both had very low nonsynonymous diversity, hinting at their likely functional importance and bringing in to question the inconclusive role of *AFB1* in auxin signaling. *AFB3* and *AFB4* had higher nonsynonymous diversity, while their sister genes, *AFB2* and *AFB5* were more conserved. This matches our current understanding of *AFB3* as playing a minor role in the auxin signaling pathway (Dharmasiri et al. 2005) and suggests *AFB4* may be undergoing pseudogenization, especially when paired with its low expression levels (Prigge et al. 2016).

**Figure 6.**
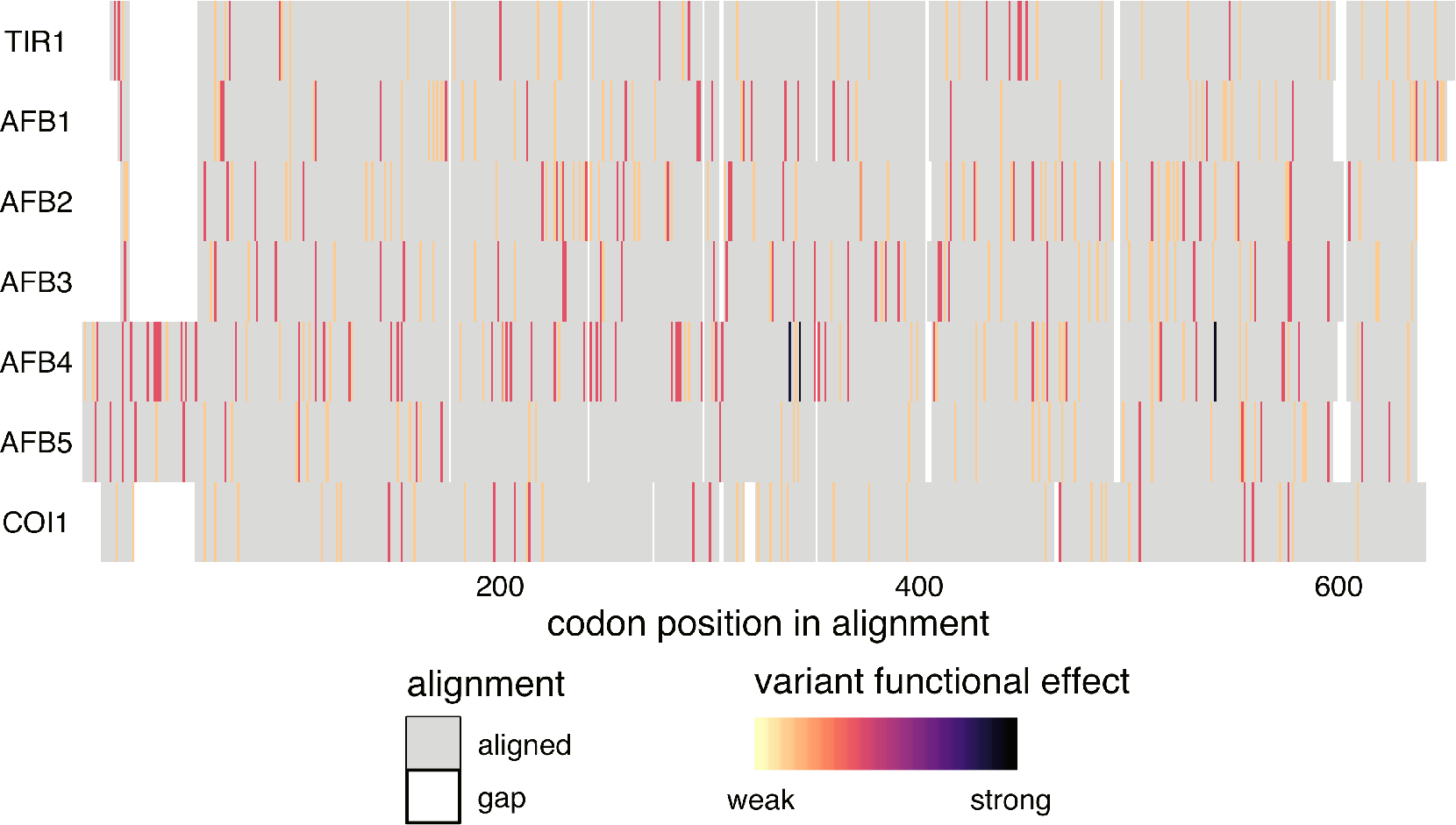
Alignment of the AFB family. Protein sequences were aligned with DECIPHER (Wright 2015) and variants were mapped to this alignment and colored according to the predicted functional effect of the allele of strongest effect at that position, with light colors having weaker effects on function and darker colors stronger effects. Red indicates missense variants. Color scale is explained in Methods.

Although most known functional regions are highly conserved in *AFB1*, there are some nonsynonymous polymorphism in the oligomerization domain that are only present in single accessions (F125E in Can-0 and I163N in Pu2-23). Mutations in this domain of *TIR1* frequently have a semidominant effect on root phenotypes (Dezfulian et al. 2016; Wright et al. 2017). Characterization of this allele and accession may help determine the role of *AFB1* in this pathway.

The AFB4 and AFB5 receptors have an N-terminal extension prior to their F-box domains. This extension had very high nonsynonymous diversity (Figure 6, suggesting that this extension does not play an important functional role in these proteins. Additionally, two frameshift variants and one stop-gained variant were observed in *AFB4* supporting its pseudogenization.

### Aux/IAA genes

For simplicity we have included only the alignment of the class A Aux/IAAs in the main manuscript. For completion we include here the complete alignment of the family.

**Figure 7.**
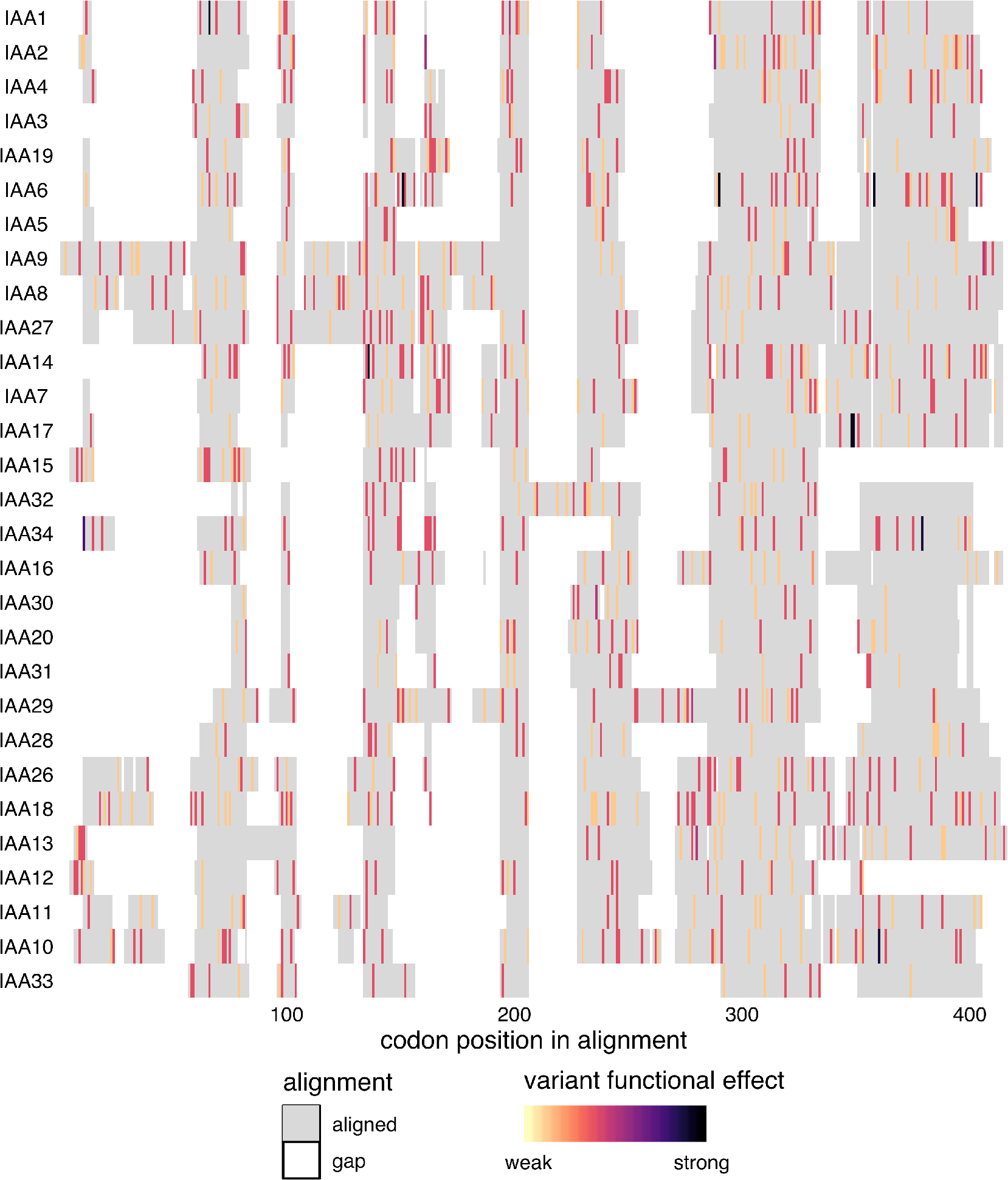
Alignment of the complete Aux/IAA family. Protein sequences were aligned with DECIPHER (Wright 2015) and variants were mapped to this alignment and colored according to the predicted functional effect of the allele of strongest effect at that position, with light colors having weaker effects on function and darker colors stronger effects. Red indicates missense variants. Color scale is explained in Methods.

**Figure 8.**
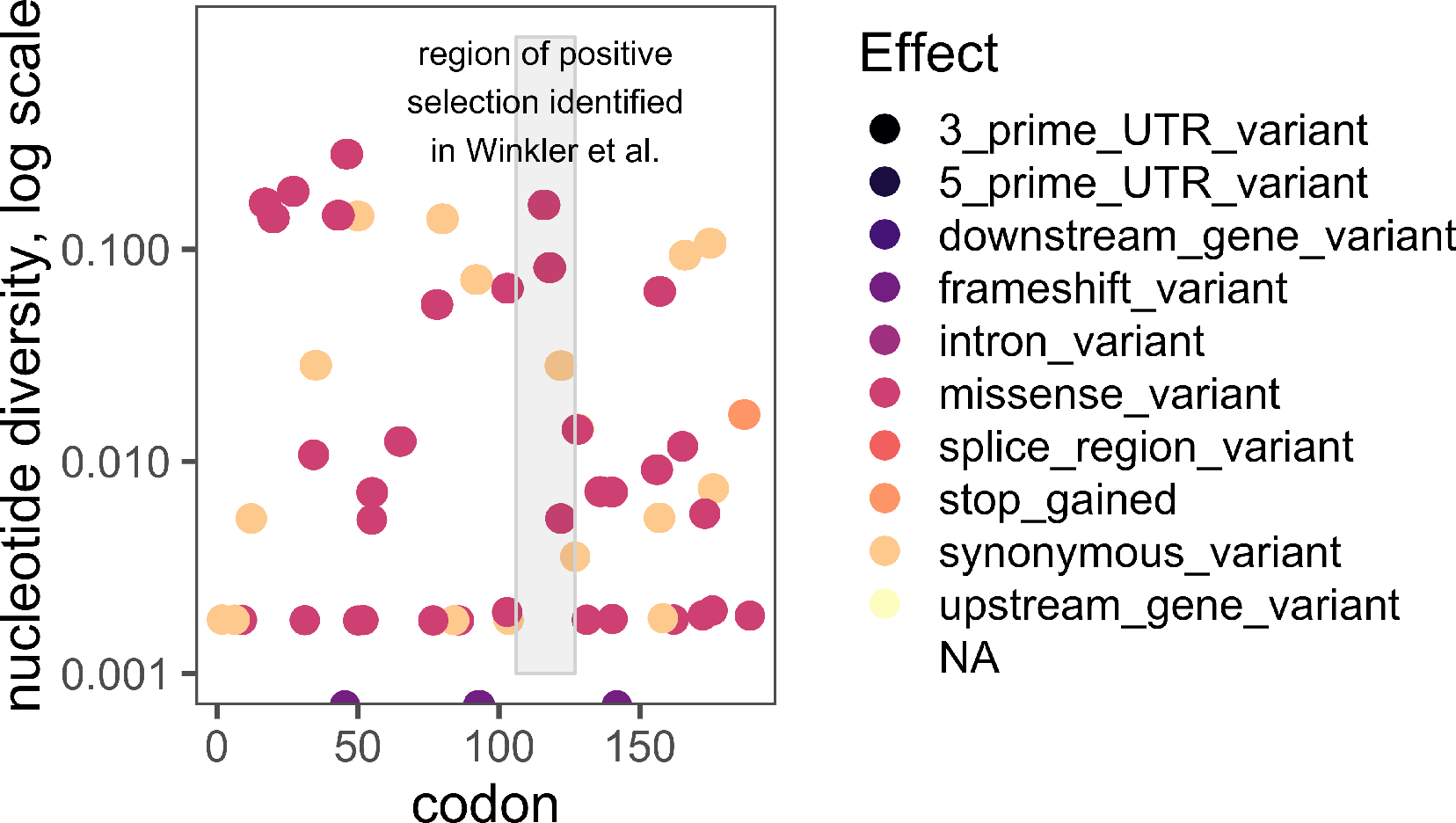
IAA6 diversity plot. Nucleotide diversity of variant positions throughout the IAA6 coding sequence are plotted and colored according to the effect of the variant alleles at each position. The region of positive selection identified by Winkler et al. is highlighed.

### TPL/TPR genes

The auxin signaling pathway utilizes the TOPLESS (TPL) and TOPLESS-related (TPR) family of Gro/TLE/TUP1 type co-repressor proteins to maintain auxin responsive genes in a transcriptionally-repressed state in the absence of auxin (Szemenyei, Hannon, and Long 2008). In *Arabidopsis thaliana* the five member *TPL*/*TPR* family includes *TPL* and *TPR1-4*. The resulting proteins are comprised of three structural domains: an N-terminal TPL domain and two WD-40 domains (Long et al. 2006). TPL/TPR proteins are recruited to the AUX/IAA proteins through interaction with the conserved Ethylene-responsive element binding factor-associated amphiphilic repression (EAR) domain (Szemenyei, Hannon, and Long 2008). Canonical EAR domains have the amino acid sequence LxLxL, as found in most AUX/IAAs (Overvoorde et al. 2005). TPL/TPR co-repressors bind EAR domains via their C-terminal to LisH (CTLH) domains found near their N-termini (citations of pre-structure founding papers/reviews). Recent structural analyses of the TPL N-terminal domain have highlighted the precise interaction interface between TPL and AUX/IAA EAR domains, as well as the TPL-TPL dimerization and tetramerization motifs (Martin-Arevalillo et al. 2017; Ke et al. 2015). The residues required for higher-order multimers of TPL tetramers have also been identified (Ma et al. 2017). Additional interactions with transcriptional regulation and chromatin modifying machinery are likely mediated by two tandem beta propeller domains of TPL/TPRs.

The TOPLESS co-repressor family generally exhibits a high level of sequence conservation at the amino acid sequence level across resequenced *Arabidopsis thaliana* accessions, with all *π*_*N*_/*π*_*S*_ values below 1 (Figure 9). The closely related *TPL* and *TPR1* have the highest *π*_*N*_/*π*_*S*_ values, suggesting that these these two related genes tolerate a higher degree of sequence and potentially functional diversity compared to *TPR2/3/4*. The N-terminal TPL domain of the TPL/TPR family is particularly conserved. All nonsynonymous polymorphisms observed in this region are either in the coils between helices or are highly conservative mutations within helices (i.e. valine to isoleucine), which would be predicted to exhibit little effect on folding and function.

**Figure 9.**
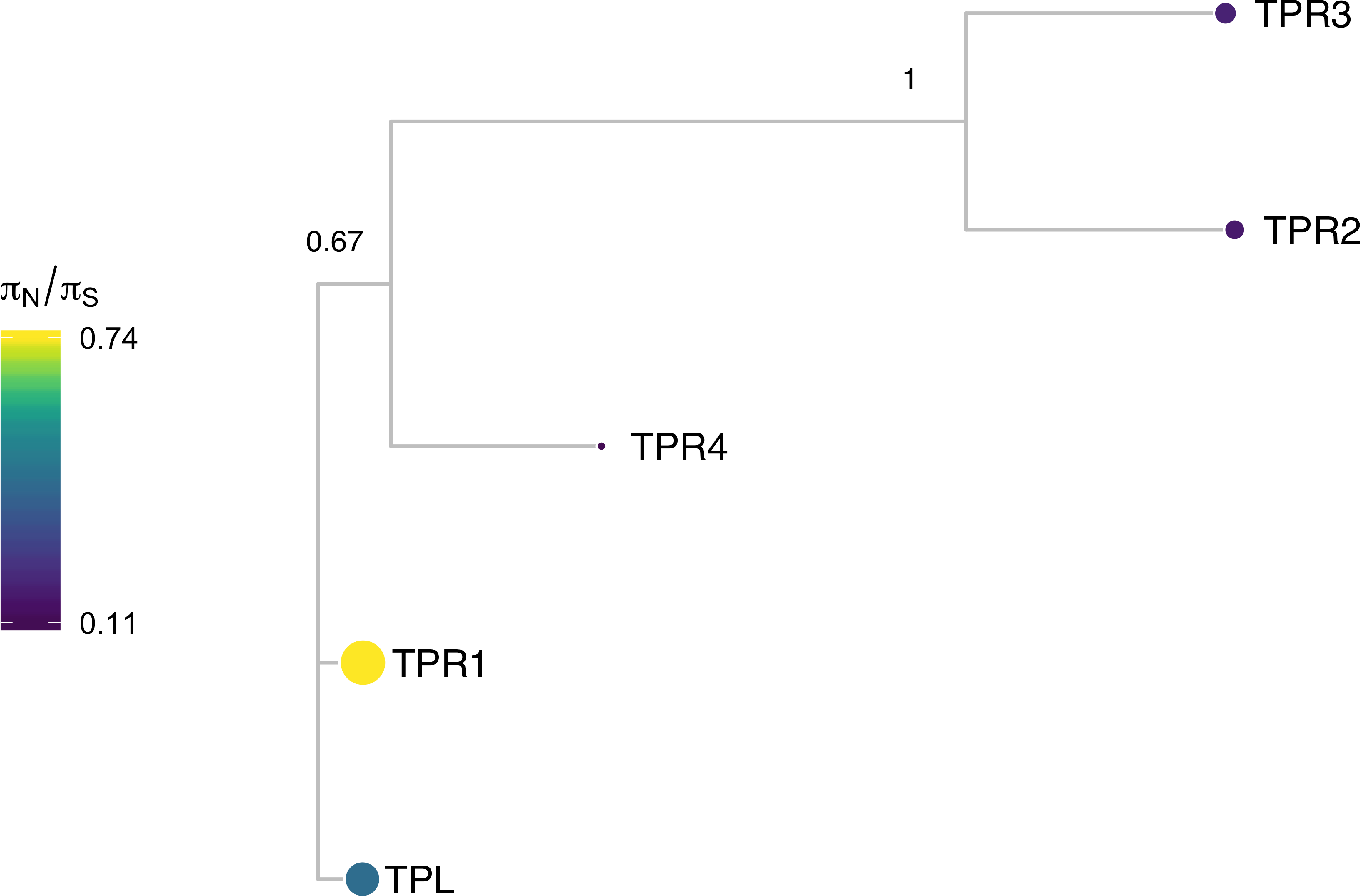
TPL protein sequence tree mapped with *π*_*N*_/*π*_*S*_. Protein sequences were aligned with DECIPHER (Wright 2015) and low information content regions were masked with Aliscore (Kück et al. 2010) prior to inferring a phylogeny with MrBayes (Ronquist and Huelsenbeck 2003). Tips of the tree are mapped with circles of diameter proportional to *π*_*N*_/*π*_*S*_ and also are colored according to *π*_*N*_/*π*_*S*_. Nodes are labeled with the poster probability of monophyly.

The high degree of conservation in the entire N-terminal domain underscores its importance in TPL/TPR function (Figure 10. For example, the initial *tpl-1* mutation (N176H) in the ninth helix is a dominant gain-of-function allele (Long et al. 2006), which is capable of binding wild-type TPL protein and inducing protein aggregation (Ma et al. 2017). It is therefore understandable that this helix had very low diversity as nonsynonymous variants in this domain could act in a dominant negative fashion.

**Figure 10.**
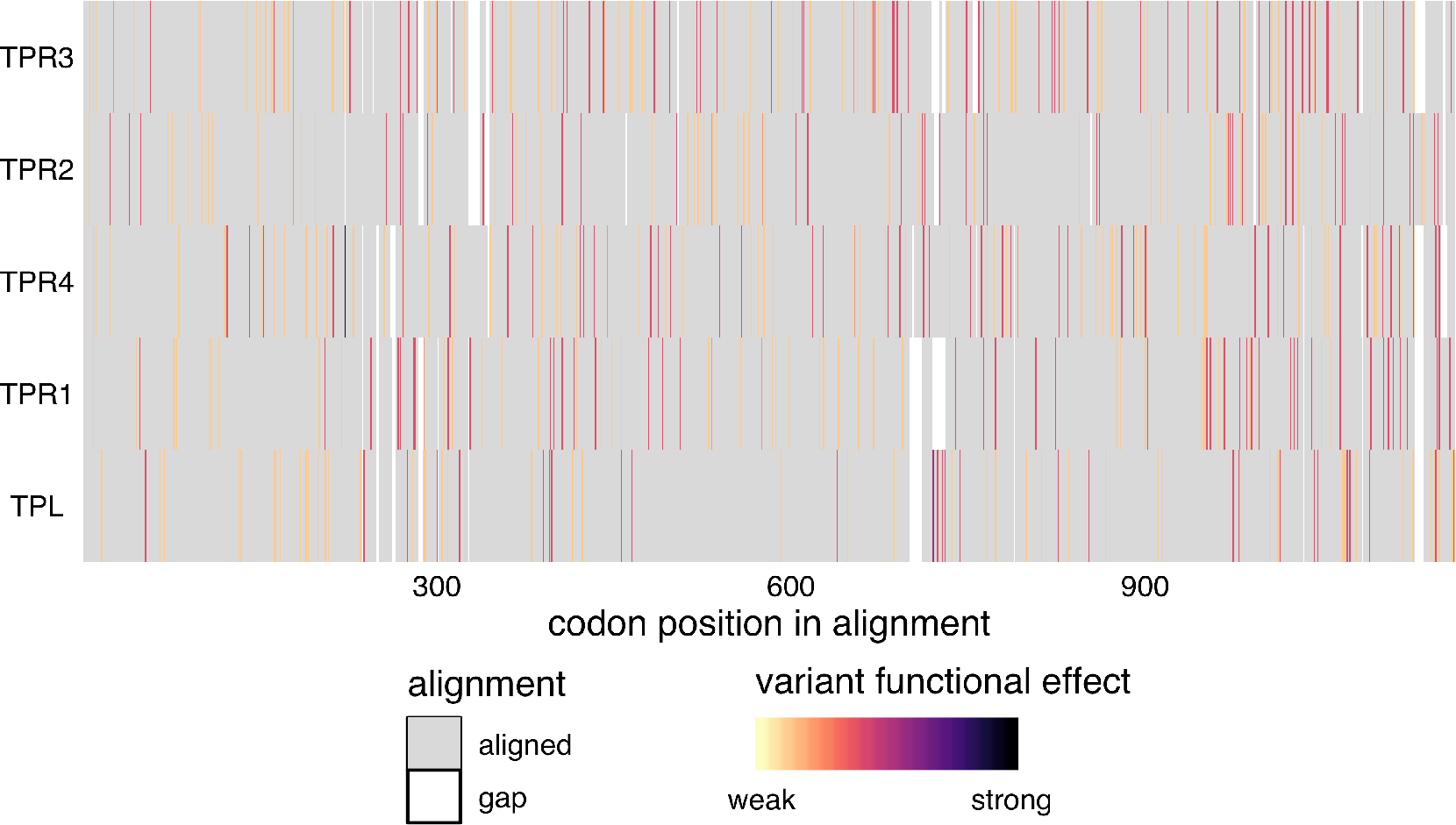
Alignment of the TPL/TPR family. Protein sequences were aligned with DECIPHER (Wright 2015) and variants were mapped to this alignment and colored according to the predicted functional effect of the allele of strongest effect at that position, with light colors having weaker effects on function and darker colors stronger effects. Red indicates missense variants. Color scale is explained in Methods.

## ARF genes

Auxin response is mediated by the auxin responsive transcription factors (ARFs). There are 23 *ARFs* in *Arabidopsis thaliana* that are divided into three phylogenetic classes. Class A ARFs (ARF5, ARF6, ARF7, ARF8 and ARF19) activate transcription. These ARFs have a glutamine-rich region in the middle of the protein that may mediate activation (Guilfoyle and Hagen 2007). It has recently been shown that the middle region of ARF5 interacts with the SWI/SNF chromatin remodeling ATPases BRAMA and SPLAYED, possibly to reduce nucleosome occupancy and allow for the recruitment of transcription machinery (Wu et al. 2015). Additionally, ARF7 interacts with Mediator subunits, directly tethering transcriptional activation machinery to its binding sites in the chromosome (Ito et al. 2016). Class B and C ARFs are historically categorized as repressor *ARFs*, though the mechanism through which they confer repression has not been identified. Their middle regions tend to be proline-and serine-rich.

Canonical ARFs are comprised of three major domains. Recent crystallization of these domains have informed structure-function analysis of the ARFs (Boer et al. 2014; Korasick et al. 2014; Nanao et al. 2014). These domains are conserved throughout land plants (Mutte et al. 2018). ARFs share an N-terminal B3 DNA binding domain. Flanking this DNA-binding domain is a dimerization domain, which folds up into a single “taco-shaped” domain to allow for dimerization between ARFs. There is an auxiliary domain that immediately follows and interacts with the dimerization domain. The middle region is the most variable between ARFs, as mentioned above, but is characterized by repetitive units of glutamine (class A), serine, or proline residues (classes B and C). The C-terminal domain of canonical ARFs is a PB1 protein-protein interaction domain mediating interactions among ARFs, between ARFs and other transcription factors, and between ARFs and the Aux/IAA repressors. This interaction domain was recently characterized as a Phox and Bem1 (PB1) domain, which is comprised of a positive and negative face with conserved basic and acidic residues, respectively (Korasick et al. 2014; Nanao et al. 2014). The dipolar nature of the PB1 domain may mediate multimerization by the pairwise interaction of these faces on different proteins as the ARF7 PB1 domain was crystallized as a multimer (Korasick et al. 2014). However, it is unclear whether ARF multimerization occurs or plays a significant role *in vivo*. Interfering with ARF dimerization in either the DNA-binding proximal dimerization domain or the PB1 domain decreases the ability of class A ARFs to activate transcription in a heterologous yeast system (Pierre-Jerome et al. 2016).

While domain architecture is broadly conserved among the ARFs, there are exceptional cases. Three ARFs do not contain a PB1 domain at all, ARF3, ARF13, and ARF17, and several more have lost the conserved acidic or basic residues in the PB1 domain, suggesting they may be reduced to a single interaction domain. Several ARFs additionally have an expanded conserved region within the DNA-binding domain, of unknown function. The majority of domain variation among ARFs occurs in the large B-class subfamily. The liverwort *Marchantia polymorpha* has a single representative ARF of each class (Flores-Sandoval, Eklund, and Bowman 2015). The expansion of these classes in flowering plants is the result of both whole genome and tandem duplication events (Remington et al. 2004). The growth of the ARF family may have allowed for the expansion of the quantity and complexity of loci regulated by the ARFs and subsequent expansion in their regulation of developmental processes (Mutte et al. 2018).

Class A ARFs are the most well-studied ARF subfamily—the five family members all act as transcriptional activators and have well-characterized, distinct developmental targets. Overall the diversity of class A *ARFs* was generally low, especially compared to the class B and C *ARFs* (Figure 11), suggesting that class A *ARFs* are central to auxin signal transduction and plant development. Analysis of class A *ARF* nonsynonymous diversity suggests that the majority of these *ARFs* are highly functionally conserved, with *π*_*N*_/*π*_*S*_ values much lower than 1 with the exception of *ARF19*, with *π*_*N*_/*π*_*S*_ value of 1.8. Comparing diversity within sister pairs, there is a similar trade-off as seen in most *IAA* sister pairs, with one sister being highly conserved and the other more divergent. *ARF19* and *ARF8* are the more divergent class A *ARFs*, with *π*_*N*_/*π*_*S*_ values at least three time those of their sisters, *ARF7* and *ARF6* respectively. This may suggest that ARF6 and ARF7 serve more essential purposes in plant development.

**Figure 11.**
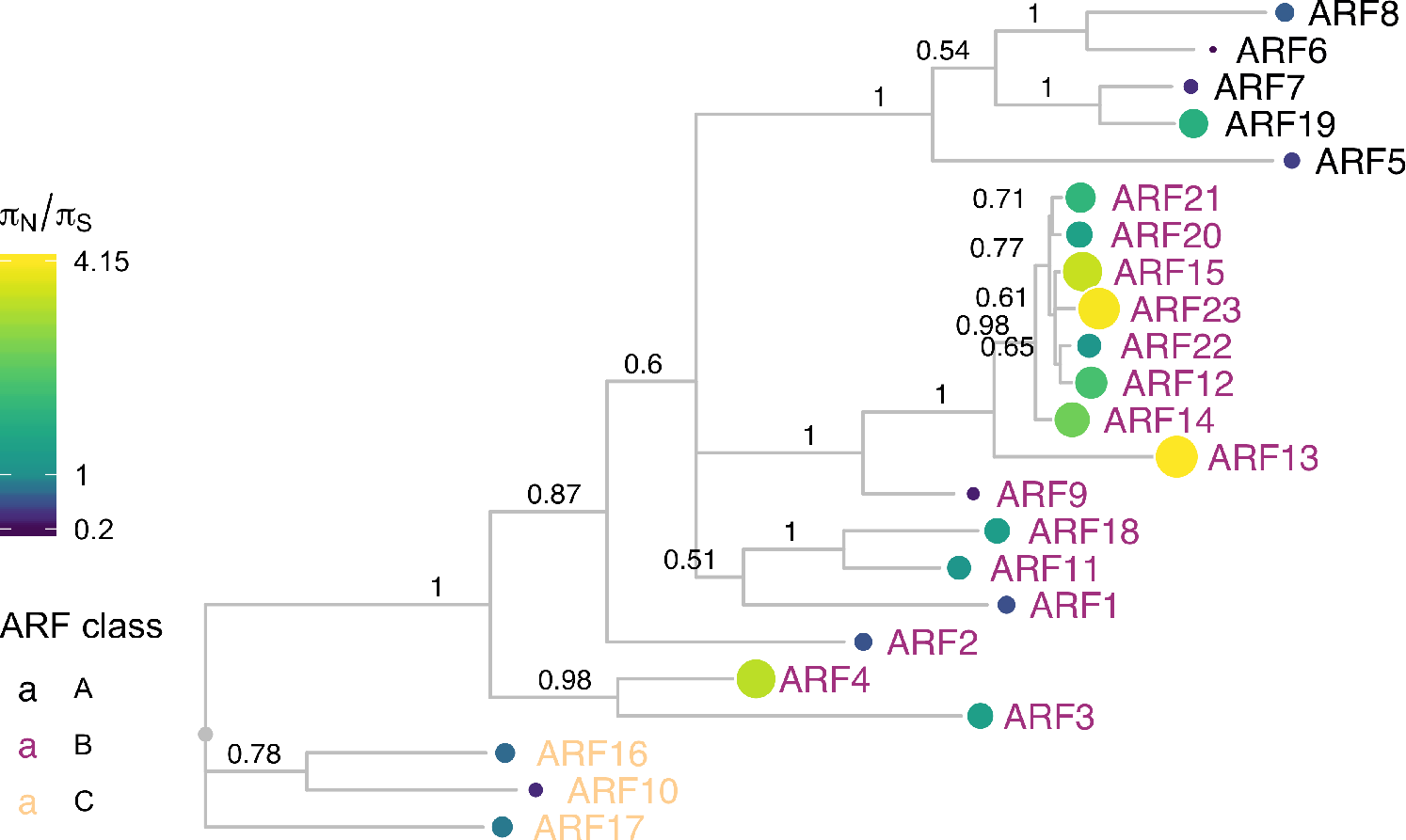
ARF protein sequence tree mapped with *π*_*N*_/*π*_*S*_. Protein sequences were aligned with DECIPHER (Wright 2015) and low information content regions were masked with Aliscore (Kuck et al. 2010) prior to inferring a phylogeny with MrBayes (Ronquist and Huelsenbeck 2003). Tips of the tree are mapped with circles of diameter proportional to *π*_*N*_/*π*_*S*_ and also are colored according to *π*_*N*_/*π*_*S*_. Nodes are labeled with the poster probability of monophyly.

For all class A ARFs, the middle region of the protein was the predominant high diversity region (Figure 12). In the analyzed natural variation, ARF7 had several expansions of polyglutamine sequences in the middle region. Polyglutamine regions are known to readily expand and contract throughout evolutionary time due to replication error, and variation in polyglutamine length can be acted on by natural selection and have phenotypic consequences (Press, Carlson, and Queitsch 2014). The ARF DNA-binding domain had very few, low-diversity missense mutations, as did the critical residues of the PB1 domain. Considering the necessity of their conserved functions, the low level of variation in these key DNA and protein-protein interaction domains is expected.

**Figure 12.**
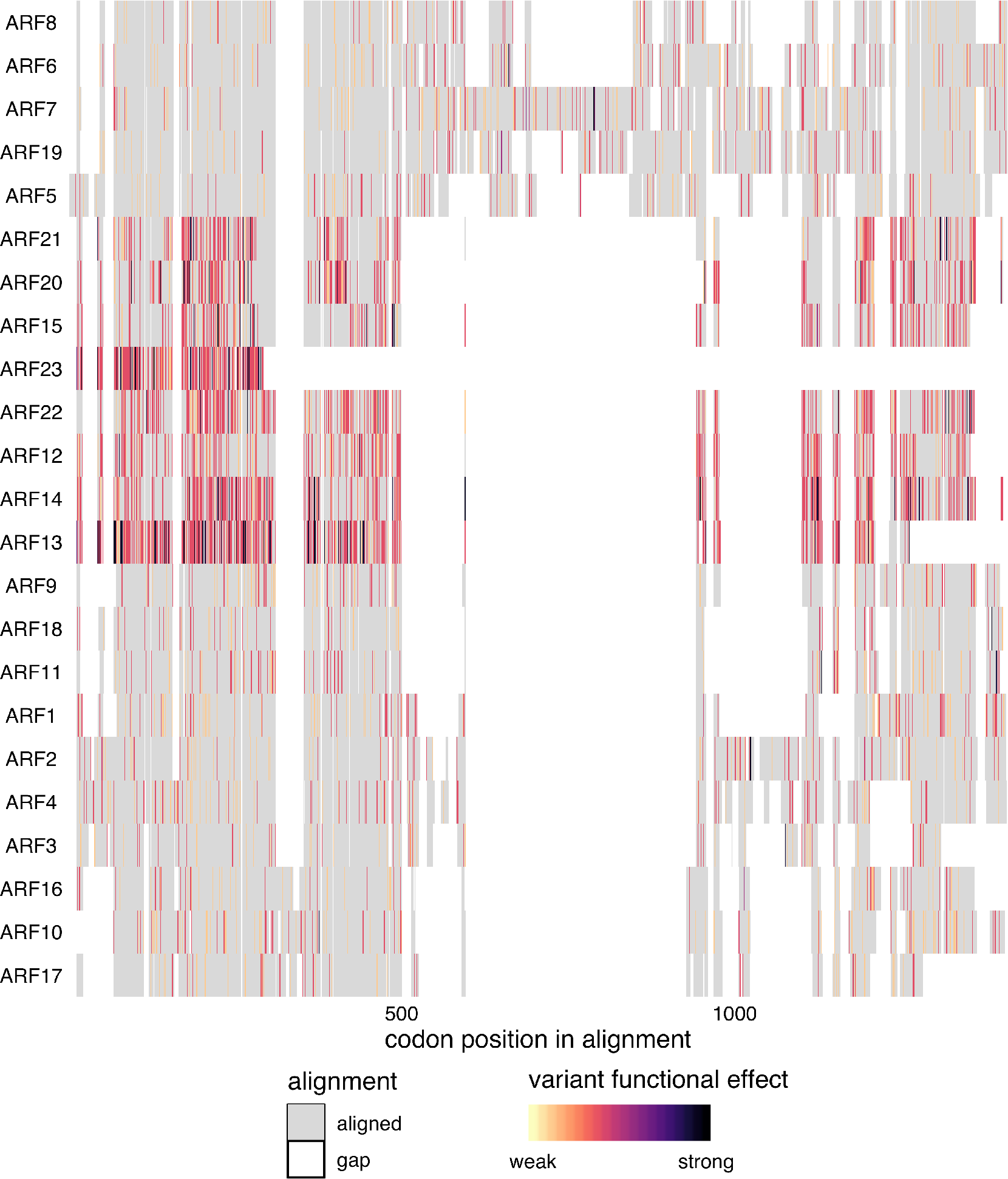
Alignment of the full ARF family. Protein sequences were aligned with DECIPHER (Wright 2015) and variants were mapped to this alignment and colored according to the predicted functional effect of the allele of strongest effect at that position, with light colors having weaker effects on function and darker colors stronger effects. Red indicates missense variants. Color scale is explained in Methods.

Many of the class B *ARFs* have very high *π*_*N*_/*π*_*S*_ ratios relative to the other *ARFs*. ARF23 has a truncated DNA-binding domain and had a high *π*_*N*_/*π*_*S*_ value of 4.1. ARF13 has many high-diversity nonsense variants and lacks a C-terminal PB1 domain. This high level of diversity, prevelance of high-frequency nonsense variants and frequent loss of critical domains, may suggest that several genes in this class are undergoing pseudogenization.

There are also a few highly conserved class B *ARFs*. The high conservation of *ARF1* and *ARF2* is expected as they play critical, redundant roles in senescence and abscission (Ellis et al. 2005). Little is known about *ARF9* however, and its low nonsynonymous diversity maybe worthy of investigation.

Class C ARFs show low nucleotide diversity scores, with all *π*_*N*_/*π*_*S*_ values substantially lower than 1. *ARF16* was the most conserved, whereas its clade members *(ARF10, ARF17)* had scores at least four times higher. Structurally, all three members of Class C ARFs contain a canonical B3 DNA-binding domain, but only ARF10 and ARF16 contain a PB1 domain. The DNA binding domains exhibit overall low diversity. Of the PB1 domain containing class C ARFs, ARF16 exhibits several missense variants which are sporadically distributed, in contrast to the conserved PB1 domain of ARF10. This conservation in the PB1 domain of ARF10 and the DBD of ARF16 may suggest subfunctionalization in this family.

